# Ecology of mosquitoes in Scottish wetlands and confirmation of *Culex molestus* and hybrids in Scotland

**DOI:** 10.64898/2026.01.20.700351

**Authors:** Georgia Kirby, Rebecca E. Brown, Meshach Lee, Jean-Philippe Parvy, Susanne Krabbendam, Emilie Pondeville, Colin Johnston, Jolyon M. Medlock, Alexander G.C. Vaux, Luca Nelli, Francesco Baldini, Heather M. Ferguson

## Abstract

**Background:** The recent spread of mosquito-borne pathogens and their vectors within Europe highlights the impact of climate change on vector-borne disease (VBD) distributions. Mosquito surveillance has been implemented in many European countries to monitor expansion of vector populations and VBDs, but ability to predict disease risk is constrained by geographic data gaps, particularly in northern areas. In the United Kingdom, wetland mosquito surveillance has been conducted extensively in England, with a knowledge deficit for Scotland. Here, we addressed this gap through a nationwide survey of mosquitoes at Scottish wetlands, with aims of 1) confirming the geographic distribution and environmental drivers of mosquito occurrence and abundance, and 2) identifying the presence of vector species of epidemiological concern, including the *Culex pipiens molestus* biotype, an important vector of emerging VBDs in mainland Europe.

**Methods:** Monthly mosquito sampling was conducted between June and October 2023 at 22 sites across Scotland comprising six wetland types: coastal saltmarsh, wet grassland, wet woodland, reedbeds, ponds and blanket bog. Adult and larval populations were sampled using Biogents BG-Pro traps and larval dipping respectively. Microclimatic and hydrological variables were recorded at collection sites and used in generalised linear mixed models to identify predictors of mosquito presence and abundance.

**Results:** 1951 adults (17 species/ groups) and 860 larvae (six species/ groups) were collected from wetlands over 183 and 164 sampling events respectively. Mosquitoes were widely distributed across the Scottish mainland including up to the northern coast, being found at all but one site. Several potential vector species including *Culex pipiens* s.l. and *Anopheles claviger* were common. Amongst the adult *Culex pipiens* s.l. specimens, approximately 8% were *Culex pipiens* biotype *molestus* or hybrid forms. Total mosquito abundance and that of key vector species were positively associated with temperature and rainfall.

**Conclusions:** We report the widespread distribution of mosquitoes in wetlands throughout Scotland, including potential vector species previously unconfirmed in Scotland. Predicted associations between mosquito abundance, rainfall and temperature indicate that climate change could favour mosquito populations in Scotland. Our results provide the first comprehensive description of mosquito ecology in Scotland, as required to update assessment of VBD risk under climate change.

## Background

Recent decades have seen the emergence of several mosquito-borne pathogens across Europe (1, 2), as well as the re-emergence of previously eliminated diseases such as malaria (3, 4). These trends are indicative of environmental changes that favour the expansion of mosquitoes and their competence to transmit pathogens. In particular, changes to climate and land use are driving disease emergence into regions previously thought to be unsuitable for transmission (1, 5). In some cases, outbreaks of vector-borne diseases (VBDs) in Europe have been attributed to the establishment of invasive mosquito species such as *Aedes albopictus*, which has been linked to autochthonous cases of dengue and chikungunya (6). However, other emerging and re-emerging diseases are spread predominantly by native mosquito populations following pathogen incursion. This includes occasional local transmission of the malaria parasite *Plasmodium vivax* by native *Anopheles* mosquitoes (4), as well as cases of zoonotic viruses such as West Nile Virus (WNV) and Usutu Virus (USUV) that are predominantly transmitted by *Culex* mosquitoes (7). Increasing numbers of WNV and USUV cases have been reported in Europe in the past two decades (8–10), especially in areas populated by the vector species *Culex modestus* and *Culex pipiens* s.l. (11).

Predicting and controlling mosquito VBD emergence requires robust mosquito surveillance. Surveillance programmes have been introduced in several European countries in recent years, particularly in southern and central Europe where *Aedes albopictus* is established and/ or WNV is endemic (12–14). Invasive mosquito surveillance (IMS) is also ongoing in parts of northern Europe (15–17), but the perceived lower risk of mosquito VBDs at higher latitudes due to cooler climates has resulted in lower prioritisation of native mosquito surveillance. The occurrence of human malaria in northern Europe historically (3, 18, 19) demonstrates that climate is not an impenetrable barrier to transmission of VBDs of human health concern in this region, with evidence indicating the risk of mosquito-borne viruses can increase rapidly with even small increases in temperature (6). Gaps in surveillance may therefore create vulnerabilities for prediction of the changing risk of VBD emergence under climate change in northern Europe, leaving countries unprepared.

Mosquito surveillance in the United Kingdom (UK) has been established in England by the government since 2010 (16, 20). Although mosquito VBD risk in the UK is likely lower than most of continental Europe due to its geographical separation and typically cooler temperatures (1, 8, 21), several mosquito species that are native to the UK are known to be competent for VBDs including WNV and USUV (22, 23) and malaria (24). The recent discovery of USUV (25–27) and WNV in England (28) demonstrates that there is a credible risk of imminent mosquito VBD establishment in the UK, while a context of historical malaria transmission by native vectors indicates that environmental conditions are not inherently unfavourable for malaria re-emergence (18). Recent surveillance indicates that 36 mosquito species occur in the UK (29), most of which are associated with wetland habitats (30). Native species with known competence for WNV include *Culex modestus, Culex torrentium, Aedes detritus* and members of the *Culex pipiens* s.l. complex (8). Amongst the latter, *Culex pipiens* biotype *molestus* (hereafter *Culex molestus*) poses a particular risk for zoonotic spillover given its inclination to bite both humans and birds (8).

Current data on mosquito distribution in the UK are largely restricted to England and Wales, where conditions are generally thought to be most climatically-suitable for mosquito-borne disease emergence (31, 32). This has resulted in a substantial knowledge gap on mosquito ecology and VBD risk in Scotland, where data are extremely sparse. Historical accounts suggest that mosquitoes have caused biting nuisance in parts of Scotland (33) and that malaria was common along the east coast and Scottish Borders in the 1700s (34). Nineteen mosquito species have been documented in Scotland from historical records, citizen science reports and sporadic field observations (35), but there has been no geographically systematic or recent survey as required to elucidate their ecology and confirm the presence of potential vector species. Notably, while the anthropophilic *Cx. molestus* biotype has been confirmed in several parts of England (36), evidence of an established presence in Scotland is unclear (35). This biotype was previously reported at a site in central Scotland between 2000 and 2003 (37), but there is no recent evidence to confirm its persistence or current distribution. Given the epidemiological risk this species poses for spillover of zoonoses from birds to humans, its investigation is warranted.

The paucity of mosquito data for Scotland constitutes a vulnerability in nationwide efforts to prepare for mosquito-borne disease emergence. Here, we aimed to address this by conducting the first comprehensive survey of adult and larval mosquitoes across Scottish wetlands. Following the model of wetland mosquito surveillance in England (30), we conducted adult trapping and larval sampling at 22 sites in Scotland encompassing five key wetland habitats with the aims of: (1) characterising mosquito species diversity, abundance and distribution, and (2) identifying environmental predictors of their abundance. We also conducted the first large-scale molecular analysis of *Cx. pipiens* s.l. mosquitoes from across Scotland, as required to assess the presence and distribution of the *Cx. molestus* biotype. In addition to generating baseline data for long-term surveillance of mosquito populations in Scotland, these results will enhance capacity to monitor and respond to mosquito-borne disease risk in the UK and northern Europe more generally.

## Methods

### Site selection

Sampling sites were selected to cover five wetland habitat types across a geographically broad area of the Scottish mainland. Target wetland habitats were ponds, reedbeds, saltmarsh, wet grassland and wet woodland (defined in **Figure 1A**), as these are known habitats for British mosquitoes that encompass all eight previously-defined ecological functional groups (**Additional file 1: Table S1**) (22), thus maximising mosquito detection and capturing species diversity. A single blanket bog site was also selected. This wetland type is limited and rare outside of Scotland, and we used this opportunity for investigation of its suitability for mosquitoes.

**Figure 1.**
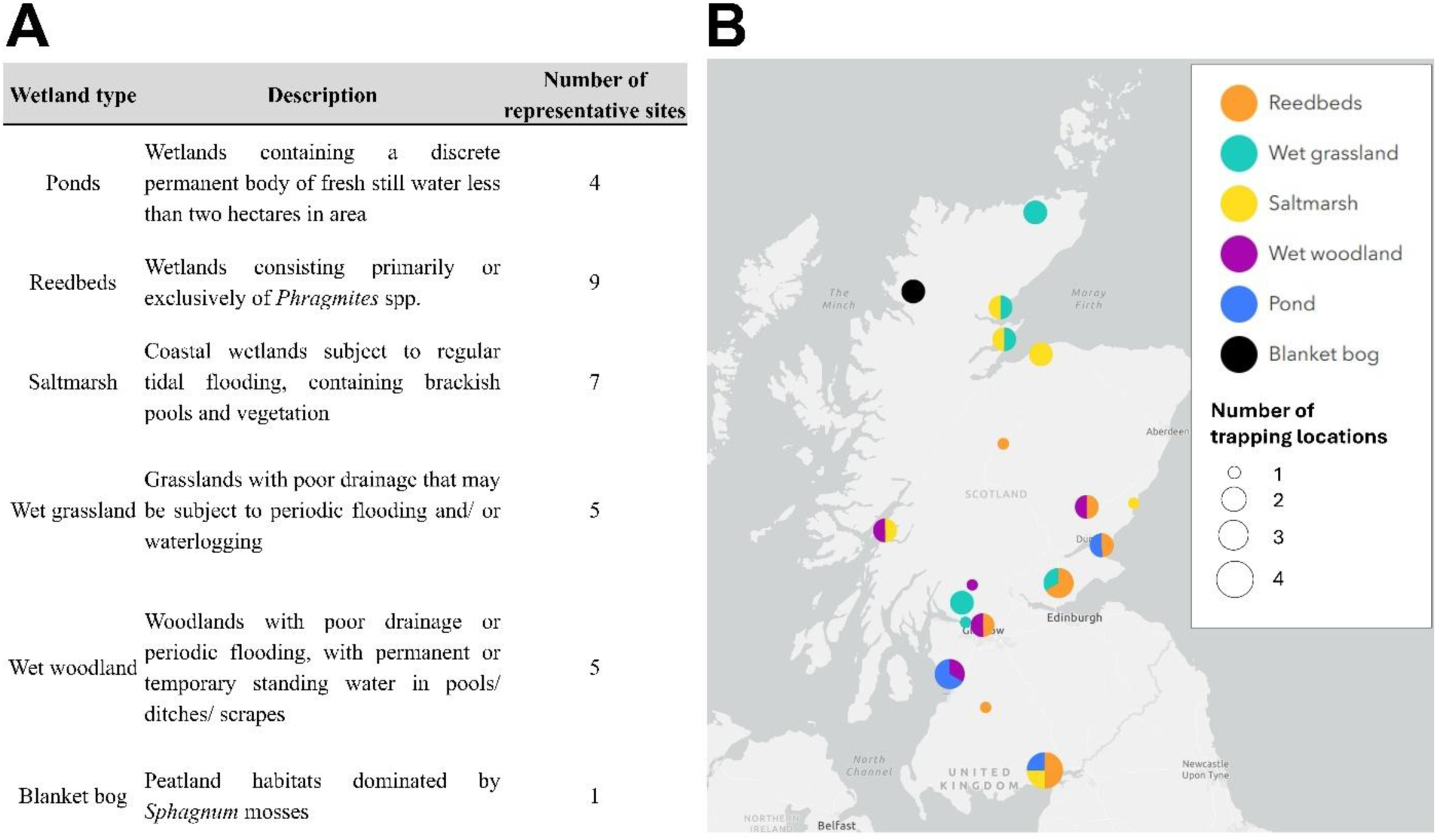
Target wetland sites. **A**. Descriptions of target wetland sites. **B**. Locations of sampling sites by wetland type. Pie charts represent the proportion of each wetland type within a 10 mile radius. Sampling sites within a 10 mile radius are aggregated.

Permission for mosquito sampling was obtained from managers of nature reserves where sites were located. We aimed to select at least one representative of each core wetland type from the northern, eastern, southern, western and central regions of the Scottish mainland. This was not always possible, but each core wetland type was represented by a minimum of four replicate sites across at least three of these geographical regions (**Figure 1B; Additional file 1: Table S2**).

### Study design

Sampling was conducted between 7th June and 4th October 2023. Where possible, sites were sampled approximately once every five weeks for adult and larval sampling. Sampling effort is detailed in **Additional file 1: Table S3**.

#### Adult sampling

One or two adult mosquito trapping locations were identified within each site. Trap locations were chosen to be within 10m of a target wetland, away from public footpaths and in sheltered locations. If reserves comprised more than one target wetland habitat, traps were placed in a maximum of two different habitat types and at least 200 metres away from each other to limit competition.

Adult trapping was conducted using battery-powered Biogents BG Pro traps (Biogents, Germany) assembled in the EVS style according to manufacturer instructions. This trap was chosen for its portability and efficiency in capturing a broad range of mosquito species (38). Traps were baited with 2kg of dry ice and a 1-octen-3-ol lure to attract a range of zoophilic species. Traps were hung from trees, bushes or fence posts no more than 1.5m from the ground. Trapping was conducted over 48-hour intervals where possible. On collection, catch bags were placed into labelled freezer bags and immediately transferred to dry ice. At one site (Findhorn Bay), mosquitoes were observed landing on members of the field team and were aspirated. Aspirated collections were placed in tubes and transferred to dry ice.

#### Larval sampling

On each sampling day, all accessible standing aquatic habitats within a 100m radius of adult traps were surveyed for larvae and pupae using a 300ml dipper, filled to approximately 250ml. Dips were made in intervals of three, with pauses of 30 seconds between each dip to allow larvae to return to the surface. The number of dips taken from each habitat within the wetland was adjusted according to the size: three dips from habitats classed as ‘small’ (< 1m^2^ in area), six dips from habitats classed as ‘medium’ (area between 1m^2^ and 3m^2^) and nine from habitats classed as ‘large’ (>3m^2^ in area). Dips were taken from at least three different areas around the edge of each habitat. Larvae and pupae were stored in labelled Falcon tubes filled with water from the habitat until processing.

#### Environmental data

GPS coordinates were recorded for all trap sites and aquatic habitats. TinyTag data loggers were attached to branches or fence posts within 1 metre of adult traps to record microclimatic temperature at 30-minute intervals. The mean of these measurements was calculated for each 48-hour trapping period. Daily temperature measurements prior to the sampling period and all daily rainfall measurements were obtained from weather station data provided by the Met Office, based on the closest weather station to each site (see **Additional file 1: Table S2**). These data were used to derive time-lagged measurements of rainfall and mean temperature up to four weeks prior to each trapping period. Time-lagged data were used to account for the effect of temperature and aquatic habitat availability during the larval development of adults caught in traps, while current microclimatic data were used to account for effects of environmental conditions on adult mosquito activity (39). Time-lagged data were calculated for the period 7-14 days prior to sampling (denoted as ‘-2’), 15-21 days prior to sampling (denoted as ‘-3’), and 22-28 days prior to sampling (denoted as ’-4’).

Hand-held meters (Hanna Instruments) and a metre stick were used to measure water temperature, pH, salinity, conductivity and depth of each aquatic habitat, with the average of three measurements recorded for each parameter. Visual observations were recorded, including the type of aquatic habitat (artificial/ ditch/ marsh/ pond/ pool/ reedbed), an estimation of the percentage of vegetation/ algal cover, and the level of exposure of the habitat (exposed/ partially shaded/ shaded). Habitat categories are defined in **Additional file 1: Table S4**. Due to lack of availability of instruments at the beginning of the sampling period and occasional malfunction of equipment, measurements were not taken for all collections.

### Mosquito processing

All adult mosquitoes and fourth-instar larvae were morphologically identified to species or species group using a British mosquito identification key (40), with additional reference to a key to European mosquitoes (41). Mosquitoes were identified under a dissecting microscope. In addition to morphology, molecular species identification was performed on adult *Culex* species due to their importance in zoonotic virus transmission (8, 42). Legs were taken from all *Culex* adults for molecular identification to distinguish morphologically identical species and biotypes. The approach taken to classify additional cryptic mosquito species groups is described in **Additional file 1: Text S1**. The species and sex were noted for each individual.

Early-stage larvae were maintained in Falcon tubes with water from the original habitat and provided with two flakes of fish food every two days until they reached the 3^rd^ and 4^th^ instar stages where morphological identification is possible. These larvae were pipetted onto a petri dish in a droplet of water and observed under a microscope for identification. No molecular identification was performed on larvae. Pupae were maintained in water-filled Falcon tubes until emergence and then identified to species as described above.

### Molecular identification of adult *Culex* specimens

PCR was conducted to distinguish morphologically cryptic *Culex* species (*Culex pipiens* s.l. from *Culex torrentium* (43), and the biotypes *Culex pipiens* biotype *pipiens* (hereafter *Culex pipiens*) and *Culex molestus* (44). Full details of this process are provided in **Additional file 1: Text S2**.

### Data analysis

All data analyses were carried out using R v. 4.5.0 (45) in R Studio (46). Generalised linear mixed models (GLMMs) were fitted using the glmmTMB package (47) to test whether environmental variables predicted mosquito abundance or presence. Global models included all biologically relevant explanatory variables (**Additional file 1: Table S5**), with site and collection date included as random effects. Latitude and longitude (derived from GPS coordinates) were included as candidate spatial predictors in global models. To reduce the risk of singularity, data were removed from sites at which only a single trapping event or collection occurred, and from the site with the sole blanket bog habitat. To test for multi-collinearity between variables, variance inflation factors (VIFs) were estimated. Variables with VIF values higher than 5 were excluded from the model (48). Continuous variables were centred and scaled to aid model fitting and facilitate comparison of effect sizes. Residual diagnostics were simulated and checked using the DHARMa package (49). Minimum adequate models were determined by backwards elimination using likelihood ratio tests. Variables were progressively removed using the lme4 *drop1* function until only those with a p-value of <0.05 remained in the model. In cases where a categorical variable was found to be a significant predictor, a post-hoc Tukey test was conducted to determine which groups differed significantly from one another. Predicted values from models were extracted using the *effects* package (50, 51). Results were plotted using *ggplot2* (52).

#### Adult abundance analysis

Four GLMMS were used to test for associations between environmental factors and the following outcomes: total mosquito abundance per trap collection (all species), and the respective abundance of *Culex pipiens* s.l., *Anopheles claviger* and *Aedes detritus*. These species were chosen due to their vector potential and relatively high prevalence. Abundance was calculated as the total number of mosquitoes per collection. Trapping periods were typically 48 hours but with some variation, so the log of number of trap nights was included as an offset. Aspirated mosquitoes were excluded from all analysis. Shannon diversity index was calculated for each trap collection using the *vegan* package (53).

Concurrent and time-lagged meteorological variables were included as fixed effects to capture the effect of microclimate on adult activity and prior larval development. Wetland type could not be fitted as a fixed effect in the model of *Aedes detritus* abundance due to a low number of observations. For each model, overdispersion was checked using the package ‘performance’. Where the data were overdispersed, negative binomial distributions were used. To determine whether linear parameterisation or quadratic parameterisation was more appropriate for each negative binomial model, global models of each were compared using Akaike Information Criterion (AIC) and the best fitting model was chosen.

#### Larval presence models

GLMMs were used to identify predictors of mosquito larval presence. Separate models were run for the two most abundant species groups: *Culex pipiens* s.l./ *Culex torrentium* (hereafter *Culex pipiens* s.l.) and *Culiseta annulata/ Culiseta alaskaensis/ Culiseta subochrea* (hereafter *Culiseta annulata*). Other species did not occur at sufficient frequencies to fit robust models. Models were fitted with a binomial distribution (0 = absent, 1 = at least one larva detected). As hydrochemical measurements were not available for all collections, two separate models were run for each species: one using the full dataset (n = 156 collections), and one using the subset included all hydrochemical measurements (n = 109 collections). Due to multi-collinearity, habitat type could not be included as a fixed effect in this model.

#### Larval density models

Separate models were used to investigate the density of two species groups: *Culex pipiens* s.l./ and *Culiseta annulata*. Larval density was defined as the total number of larvae collected from a habitat on a given date, with the log of the number of dips taken from the habitat included as an offset. GLMMS to identify predictors of larval density were fitted with a negative binomial regression distribution due to overdispersion in the response variable. Due to low sample sizes, models were simplified to enable convergence and hydrochemical variables were not included.

## Results

### Conditions during sampling period

Details on climatic conditions are presented in **Additional file 1: Text S3** with a summary by site in **Table S6**.

### Adult mosquito populations

A total of 1,951 adult mosquitoes was collected during 95 trapping events (183 trap nights) between June and October 2023. An additional four mosquitoes were collected by aspiration at Findhorn Bay. The majority of collected mosquitoes were females (97.5%). Mosquitoes were captured at 21 of the 22 wetland sites; the only site where none were captured was the sole blanket bog site (Knockan Crag).

#### Adult species composition and distribution

Seventeen mosquito species/ species groups were sampled (**Table 1**), encompassing all eight British mosquito functional groups (30). The most abundant species was *Coquillettidia richiardii* (35.3%; **Table 1**). Mosquitoes aspirated at Findhorn Bay were *Aedes cantans* (n = 3) and *Ae. detritus* (n = 1). Amongst the specimens morphologically identified as *Culex pipiens* s.l./ *Culex torrentium*, 163 (82%) were identified by PCR as *Culex pipiens*, 10 (5.1%) were *Culex molestus,* and five (2.5%) were *Culex pipiens* x *Culex molestus* hybrids. No specimens were identified as *Culex torrentium*. A further 20 specimens (10.2%) could not be conclusively identified. *Culex molestus* and hybrids were found at three sites, where they co-occurred: Caerlaverock NNR (n = 7), WWT Caerlaverock (n = 6) and RSPB Loch Lomond (n = 2).

**Table 1.**
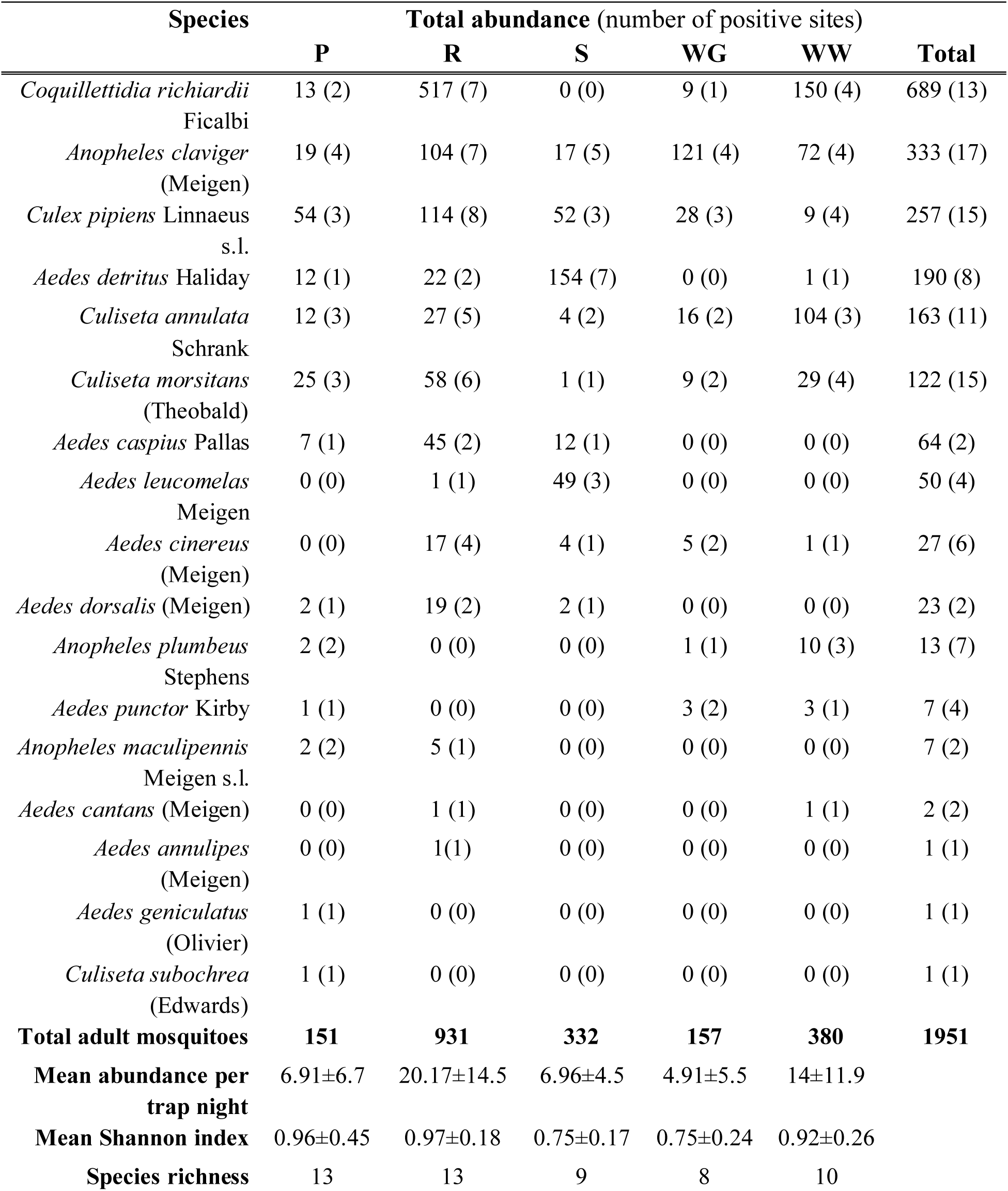
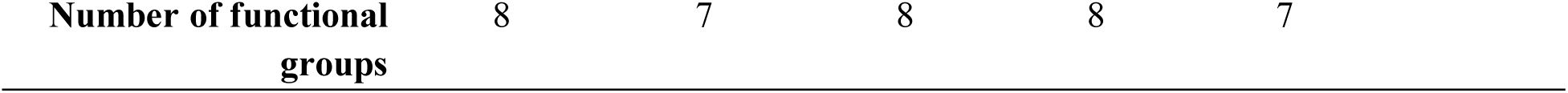
Adult mosquito species abundance in traps and presence in different wetland-associated habitats. P = Ponds, R = Reedbeds, S = Saltmarsh, WG = Wet grassland, WW = Wet woodland. Mean values include ± 95% confidence interval.

*Anopheles claviger* was the most widely distributed and ubiquitous species, occurring at 17 of 22 sites (**Table 1**; **Figure 2A**) and across all target wetland types. Three additional species (*Culex pipiens* s.l., *Culiseta annulata* and *Culiseta morsitans*) were also found in all target wetland types, but in a lower number of sites overall (**Figure 2B**). All wetland types were populated by a variety of species representing all, or almost all, functional groups (**Additional file 1: Figure S1**), with species richness and diversity generally highest in pond and reedbed sites.

**Figure 2.**
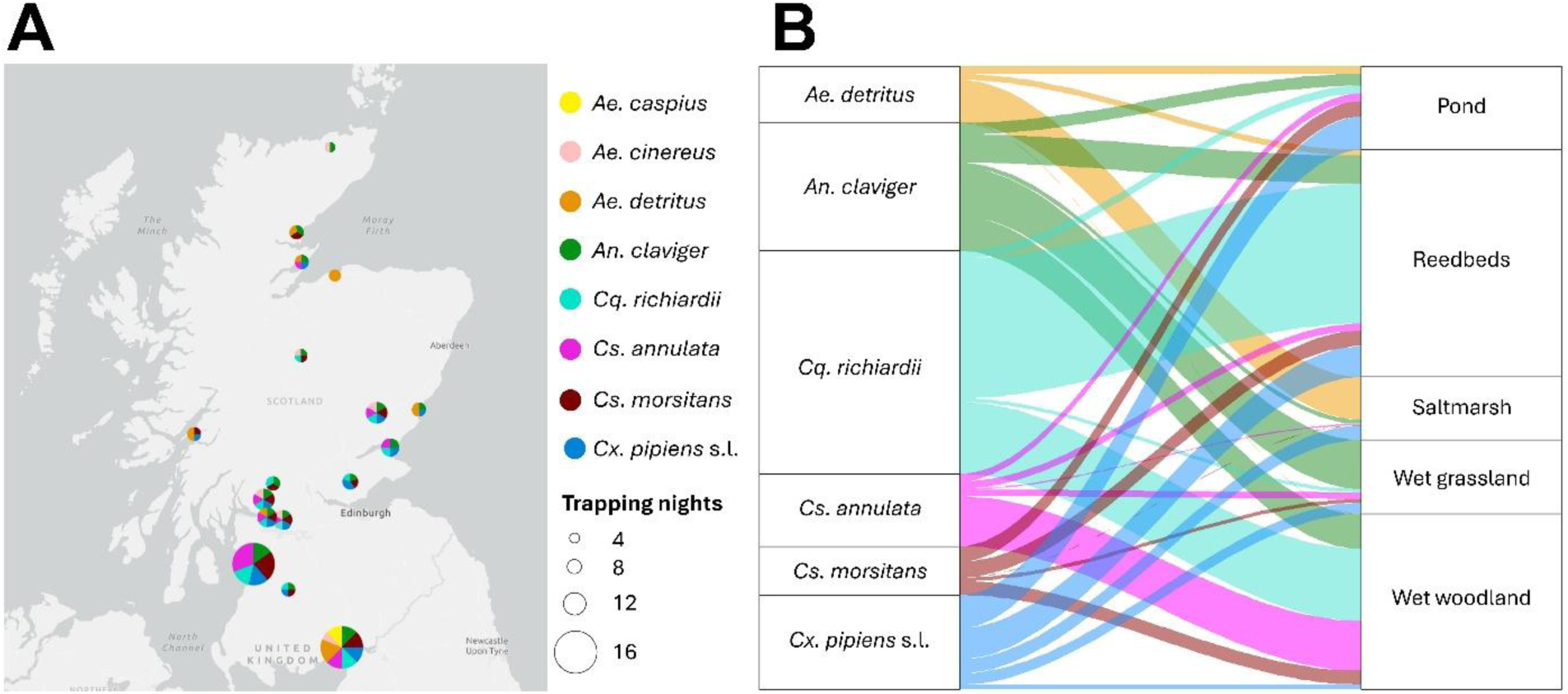
Distribution and associated wetland habitats of the most abundant adult mosquito species. **A.** Distribution of adult-stage species throughout mainland Scotland. Trap sites within a 10-mile radius are aggregated. **B.** Allocation of common adult-stage species by wetland type. Ribbon width indicates abundance.

#### Predictors of adult abundance

The study was designed to maximise spatial coverage over temporal sampling, with sites sampled too infrequently for detailed elucidation of seasonality at each site. However, there was a trend of higher mosquito numbers in collections made between July and early August – partly due to high abundance of *Coquillettidia richiardii* at reedbed sites during this period (**Additional file 1: Figure S2**). Adult mosquito abundance was positively associated with temperature and rainfall in both total mosquito and species-specific analyses. Mean temperature during the trapping period was positively associated with total mosquito (χ^2^ = 5.344, p = 0.021; **Figure 3A**) and *Culex pipiens* s.l. abundance (χ^2^ = 3.941, p = 0.047; **Figure 3B**). The abundance of *Anopheles claviger* was positively associated with rainfall during the trapping period (χ^2^ = 4.017, p = 0.045; **Figure 3C**) and three weeks prior to trapping (χ^2^ = 7.327, p = 0.007; **Figure 3D**). Lagged rainfall was also positively associated with total mosquito abundance (three weeks before collection; χ^2^ = 6.304, p = 0.012; **Figure 3E**) and *Aedes detritus* (four weeks before collection; χ^2^ = 14.263, p = <0.001; **Figure 3F**).

**Figure 3.**
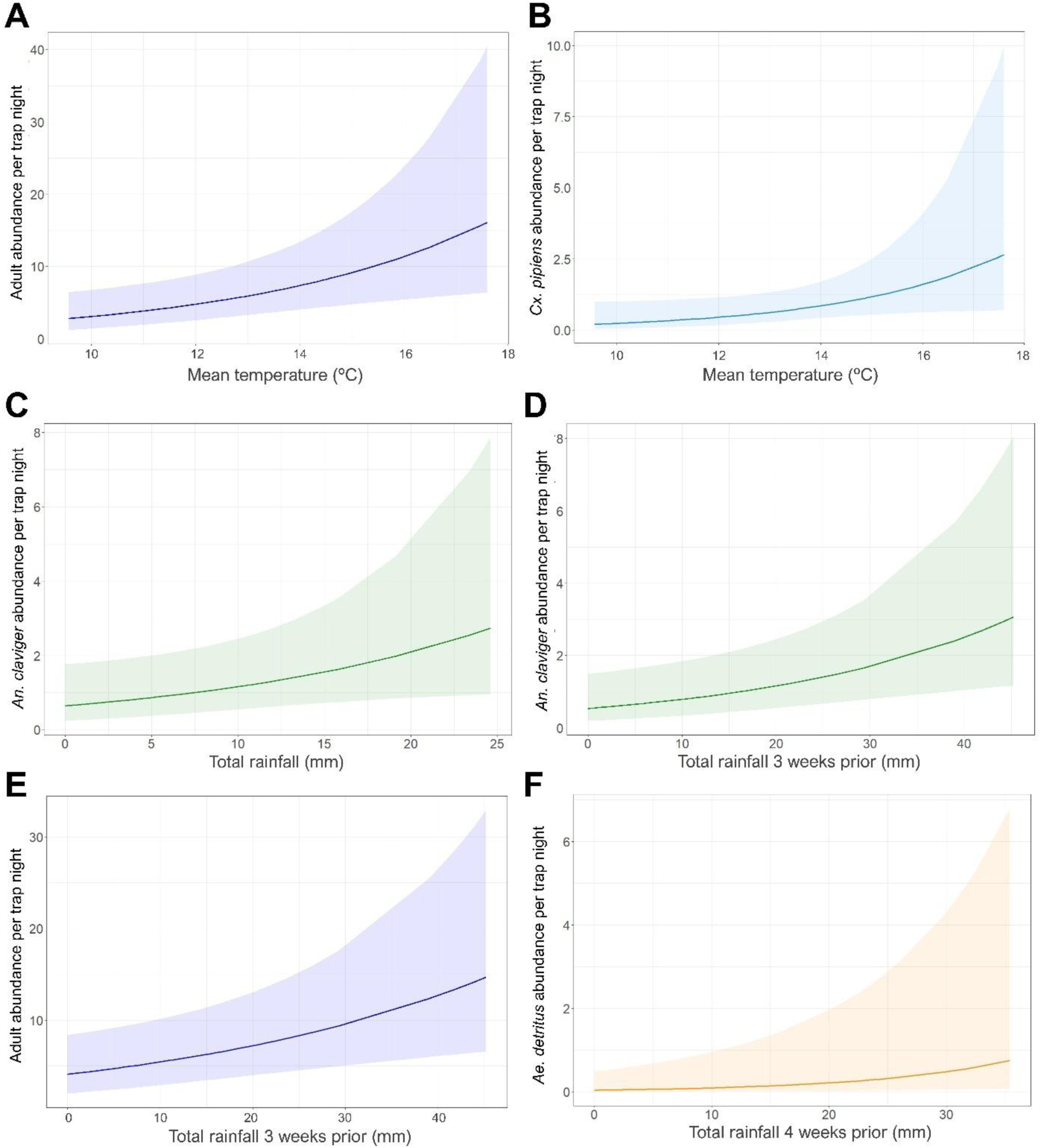
Environmental predictors of adult mosquito abundance. Shaded areas indicate 95% confidence intervals. **A.** Predicted relationship between concurrent mean temperature and total adult abundance. **B.** Predicted relationship between concurrent mean temperature and *Culex pipiens* s.l. abundance. **C.** Predicted relationship between concurrent rainfall and *Anopheles claviger* abundance. **D.** Predicted relationship between rainfall three weeks prior to sampling and *Anopheles claviger* abundance. **E.** Predicted relationship between rainfall three weeks prior to sampling and total adult abundance. **F.** Predicted relationship between rainfall four weeks prior to sampling and *Aedes detritus* abundance.

Latitude was negatively associated with *Cx. pipiens* s.l. abundance (χ^2^ = 4.551, p = 0.033; **Figure 4A**), while longitude positively correlated with abundance of *An. claviger*, indicating an easterly distribution (χ^2^ = 5.628, p = 0.018; **Figure 4B**). No other environmental variables, including wetland type, were significantly associated with any mosquito outcome variables (**Additional file 1: Table S7**).

**Figure 4.**
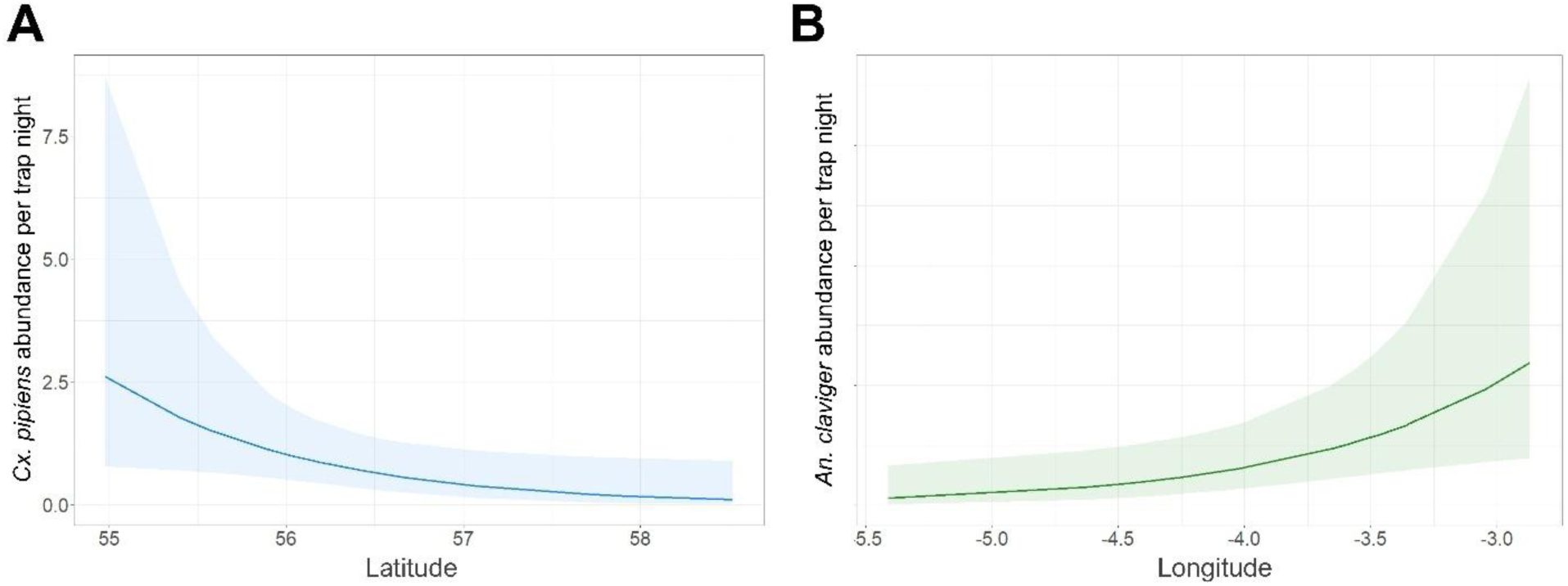
Spatial predictors of adult mosquito abundance. Shaded areas indicate 95% confidence intervals. **A.** Predicted relationship between latitude and *Culex pipiens* s.l. abundance. **B.** Predicted relationship between longitude and *Anopheles claviger* abundance.

### Larval populations

Larvae were detected in 63 of 164 sampling events from wetland habitats (38.4%). Eight hundred and sixty larvae were collected. Six species were found, the vast majority of which were *Culex pipiens* s.l. and *Culiseta annulata,* which made up 42.2% and 41.7% of collections respectively. The remaining species were *Anopheles claviger* (9.2%), *Culiseta morsitans* (4.4%), *Aedes detritus* (1.7%) and *Aedes leucomelas* (0.3%).

Larvae were found in a variety of natural and artificial habitat types (**Table 2**). Artificial habitats included bird baths, feeding troughs and artificial ponds, and were dominated by *Cx. pipiens* s.l.. *Culex pipiens* s.l. and *Cs. annulata* co-occurred in 2.4% of all collections, but only in natural habitats. Typical habitat characteristics by species are summarised in **Additional file 1: Table S8**.

**Table 2.** Breakdown of larval species by habitat types in aquatic surveys where larvae were detected.

| Species | Number of sampling events by habitat type in which larvae were detected |  |  |  |  |  |  | Total |
| --- | --- | --- | --- | --- | --- | --- | --- | --- |
|  | Artificial | Bog | Ditch | Marsh | Pond | Pool | Reedbed |  |
| *Culex pipiens* s.l. | 10 | 0 | 2 | 2 | 2 | 3 | 0 | 19 |
| *Culiseta annulata* | 3 | 0 | 5 | 4 | 2 | 9 | 5 | 28 |
| *Anopheles claviger* | 2 | 0 | 1 | 1 | 1 | 1 | 7 | 13 |
| *Culiseta morsitans* | 0 | 0 | 0 | 1 | 1 | 2 | 2 | 6 |
| *Aedes detritus/ Aedes leucomelas* | 0 | 0 | 0 | 1 | 0 | 2 | 0 | 3 |

#### Predictors of larval presence and density

No significant predictors of *Culex pipiens* s.l. larval presence were identified in analysis of the full dataset that included only macrohabitat characteristics. However, analysis of the subset of data (n =120) that included hydrochemical measurements identified a positive association between *Cx. pipiens* s.l. larval presence and water temperature (χ^2^ = 8.183, p = 0.004; **Figure 5A**) and vegetation cover (χ^2^ = 4.423, p = 0.035; **Figure 5B**). Vegetation cover was also positively associated with *Cs. annulata* larval presence in both complete and subsetted models (χ^2^ = 4.294, p = 0.038; **Figure 5C**), as was habitat exposure (χ^2^ = 8.756, p = 0.013; **Figure 5D**). Specifically, shaded habitats had a significantly higher likelihood of *Cs. annulata* presence than partially shaded and exposed habitats. Conductivity was negatively associated with *Cs. annulata* presence in the subsetted model (χ^2^ = 4.265, p = 0.039; **Figure 5E**).

**Figure 5.**
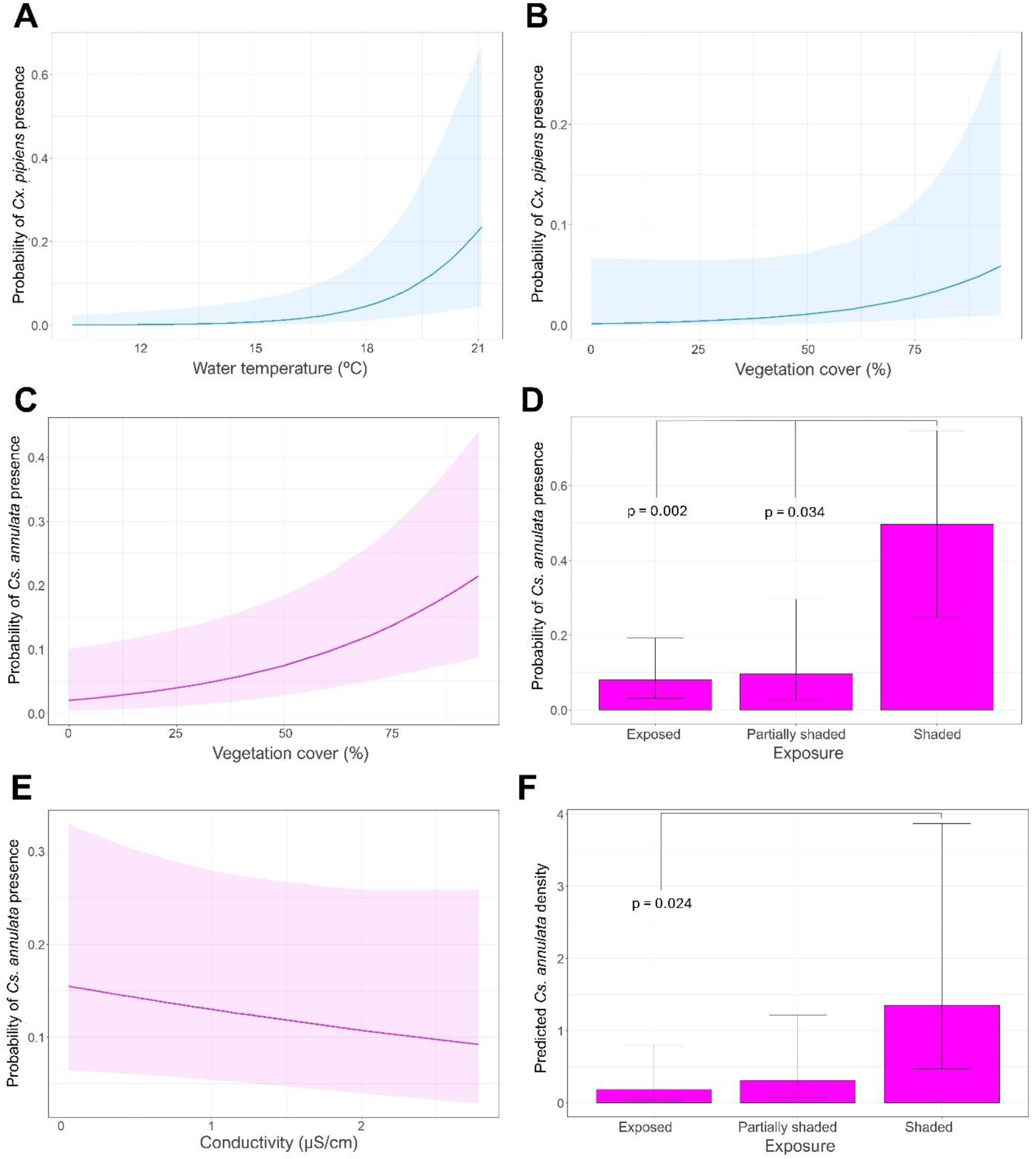
Variables associated with larval presence and density. Error bars and shaded areas indicate 95% confidence intervals. **A.** Predicted relationship between water temperature and *Culex pipiens* s.l. presence. **B.** Predicted relationship between vegetation cover and *Culex pipiens* s.l. presence. **C.** Predicted relationship between vegetation cover and *Culiseta annulata* presence. **D.** Predicted relationship between habitat exposure and *Culiseta annulata* presence. **E.** Predicted relationship between conductivity and *Culiseta annulata* presence. **F.** Predicted relationship between habitat exposure and *Culiseta annulata* density.

*Culex pipiens* s.l. and *Cs. annulata* larvae were typically found in higher densities than other species. No statistically significant predictors of *Cx. pipiens* s.l. density were detected (**Additional file 1: Table S9**). However, habitat exposure impacted *Cs. annulata* density (χ^2^ = 11.113, p = 0.004; **Figure 5F**), with shaded habitats tending to have higher larval densities than exposed habitats (p = 0.024).

## Discussion

Here we investigated the abundance and distribution of mosquitoes in Scotland, with the aim of addressing geographic data gaps that hinder assessment of mosquito VBD risk for the UK as a whole. Mosquitoes were widespread in wetlands across the Scottish mainland, with several of the species detected known to pose a risk as vectors in England and other parts of Europe. This includes *Culex pipiens* s.l., with molecular confirmation of *Culex molestus* and hybrids at several sites. Temperature and rainfall were positively associated with the abundance of several adult mosquito species, including *Cx. pipiens* s.l. These results indicate that mosquitoes are common in Scottish wetland ecosystems, and that the warmer and wetter conditions expected with climate change could increase the abundance and distribution of potential vectors.

Seventeen mosquito species were collected through adult trapping, encompassing fourteen of the nineteen species that have previously been recorded in Scotland (35) and providing new records for *Aedes dorsalis, Aedes leucomelas* and *Aedes geniculatus*. Among those species previously recorded in Scotland but absent here were *Culiseta alaskaensis* (which we acknowledge may have been present in our collections but not distinguished from the morphologically similar *Culiseta annulata*), *Culex torrentium* (which similarly may have been present in larval collections but not distinguished from *Cx. pipiens* s.l.), *Aedes rusticus*, *Aedes sticticus*, *Culex territans* and *Culiseta fumipennis*. With the exception of *Ae. rusticus* and *Cs. fumipennis,* these species are thought to be uncommon and are rarely recorded elsewhere in the UK (54). While *Cx. torrentium* is thought to be more common at northern latitudes (55), our sampling methods may have caused adults to go undetected due to a previously-documented trapping bias (56). Notably, the most commonly collected adult species here, *Coquillettidia richiardii*, has only been recorded in Scotland once previously (35). This species is ubiquitous in southern England (29), but we hypothesise that its widespread distribution in Scotland reflects the lack of previous research rather than a recent range expansion. Its vector competence is unknown at present, but populations should be monitored given their propensity to feed on both birds and humans and consequent potential to act as bridge vectors (8, 54).

At least six other native mosquito species with potential to act as vectors of zoonotic viruses and/ or human malaria were detected: *Cx. pipiens* s.l.*, Aedes caspius, Aedes detritus, Anopheles maculipennis* s.l., *Anopheles plumbeus* and *Anopheles claviger* (54, 57). Of particular note was evidence of *Cx. molestus* populations and the first discovery of *Cx. pipiens* x *Cx. molestus* hybrids in Scotland. Previously, there has been reference to a historical record of *Cx. molestus* in eastern Scotland (35) and reports of *Cx. molestus* from a seemingly isolated population in Menstrie (central Scotland) between 2000 and 2003 (37, 44, 58, 59), but to our knowledge there have been no formal records of wider *Cx. molestus* distribution or persistence in Scotland since. Results here indicate that *Cx. molestus* populations are established in wetlands in west and southern Scotland. Although northern European populations of *Cx. molestus* were initially linked to urban subterranean habitats (8, 60), recent studies confirm their presence at low levels in farmland and rural sites in England (36, 61, 62), consistent with our findings. Our detection of *Cx. pipiens* x *Cx. molestus* hybrids indicates spatial overlap of these forms in Scotland, countering previous suggestions that hybridisation is unlikely at northern latitudes due to habitat and behaviour differences (63). Given that *Cx. molestus* and hybrids are highly competent for WNV and USUV (64–66) and can be key bridge vectors due to their inclination to feed on humans and birds (67), our findings have implications for identifying zoonotic VBD risk in Scotland.

We identified several environmental predictors of adult mosquito abundance that could influence VBD risk under climate change. In line with studies from other temperate settings (68), mean ambient temperature during the trapping period was positively associated both with total adult abundance and with abundance of *Cx. pipiens* s.l. specifically, suggesting a positive effect of temperature on mosquito host-seeking activity. Average summer temperatures in Scotland are predicted to rise by 3.1°C over the next sixty years under climate change (69), which our findings here suggest could boost mosquito-host contact. Activity of *An. claviger* also appeared to increase with precipitation, while lagged rainfall (three to four weeks prior to collection) was positively correlated with total adult mosquito abundance and that of *An. claviger* and *Ae. detritus*. This likely reflects increased larval habitat availability for these and other mosquito species that can exploit transient or semi-permanent habitats such as flooded grassland and saltmarsh (5, 57, 70). The observed relationships of *An. claviger* and *Cx. pipiens* s.l. abundance with longitude and latitude respectively may be similarly indicative of habitat suitability and availability, as grassland and arable land (favouring *An. claviger*) is most prevalent in eastern regions of Scotland, while built-up areas and urban greenspaces (favouring *Cx. pipiens* s.l.) are scarce in northern parts of the country (71). Together, these findings suggest that VBD risk across Scotland is spatially heterogeneous, with microclimate and land use likely to play key roles in vector distribution, abundance and activity.

Larvae of the six most abundant adult species were found during surveillance, with the exception of *Cq. richiardii.* Failure to detect *Cq. richardii* larvae is unsurprising given their unusual larval ecology within the stems of emergent vegetation, making them highly unlikely to be found using traditional larval sampling methods (8). The ubiquity of *Cx. pipiens* s.l. and *Cs. annulata* larvae and the broad range of habitats that they were found in demonstrates the versatility of these species in Scotland, in line with what has been recorded elsewhere (72, 73). Presence of *Cx. pipiens* s.l. larvae was positively associated with water temperature. As this species group was prevalent in artificial habitats, which tend to be shallow and more exposed, this effect may have been confounded by habitat type, which could not be included in models due to collinearity. Although higher temperatures are associated with increased mortality (74), the temperatures at which *Cx. pipiens* s.l. larvae were found here (13.5 °C – 21.4°C) fall within the range for optimal survival (74, 75). However, our findings offer some insight into potential measures to offset climate change-mediated increases in *Cx. pipiens* s.l. abundance and corresponding VBD risk in Scotland; the frequency at which *Cx. pipiens* s.l. larvae were encountered in artificial habitats suggests that reduction and monitoring of such habitats could be an effective form of control in built-up areas. Despite displaying opportunistic breeding site choices, *Cs. annulata* was found relatively infrequently in artificial container habitats and never in containers occupied by *Cx. pipiens* s.l., suggesting discrepancies in habitat preference and a possible competition effect that could be further explored.

While the findings of this study significantly expand and update our knowledge of mosquito vector ecology in Scotland, limitations in the methodology may have led to underrepresentation of certain species. Uneven geographic distribution of sampling sites and wetland types, as well as a late start to sampling relative to the start of the mosquito activity season (22, 30), may have caused uncommon or univoltine (one generation per year) species and/ or the peaks of their abundance to be missed. In particular, we note low abundance of vernal human-biting woodland floodwater species such as *Aedes cantans* and *Aedes punctor*, which have been observed in high numbers in wet woodland sites in England (22) but may have been underrepresented here due to a lack of spring sampling and a disproportionate number of wet woodland sampling sites located in the west of Scotland, where mosquito densities were generally lower. We also acknowledge that some species-wetland associations may have been obscured by the high proportion of mixed-habitat sites and heterogeneity of aquatic habitats within wetland types, which may partly explain why wetland type was not a significant predictor of species abundance. Caution should therefore be taken when inferring species-wetland associations. Future surveillance should begin earlier in the season and encompass a more even spread of wetland types across the country to fully capture species diversity. Additionally, more in-depth longitudinal surveillance at fixed sites will be required to clearly describe the pattern of seasonality and inter-annual variation in mosquito abundance and human biting risk in Scotland.

## Conclusions

This study represents the first systematic survey of mosquito species diversity in Scotland, filling a geographic gap in British mosquito surveillance. Our findings show that several potential vector species are distributed throughout the mainland, including localised presence of *Culex molestus* and hybrid forms, the latter of which were detected here for the first time in Scotland. Microclimatic variables were found to positively correlate with Scottish mosquito abundance and activity, suggesting that climate change-mediated increases in spring/ summer temperature and rainfall events could lead to higher numbers of potential vector species in Scottish wetlands. The findings here have relevance for other northern regions of Europe and should be incorporated into ecological and epidemiological models to evaluate the risk of VBD risk at northern latitudes under climate change.

## Declarations

## Acknowledgements

We are grateful to RSPB Scotland, the Scottish Wildlife Trust, the Wildfowl and Wetlands Trust, Forestry and Land Scotland, West Dunbartonshire Council and Moray Council for site access and permissions. We are also grateful to Nina Gangi for contribution to pilot studies of this work and to Colin Malcolm for useful discussions about *Culex molestus*. We thank Sarah Kelly and members of the UK VBD Hub team for assistance with data curation and deposit.

## Availability of data

The dataset used in this study was deposited at VecDyn through the VBD Hub with curation support from Sarah Kelly, and is available for download by request.

## Author contributions

H.M.F., E.P., J.M.M, A.G.C.V., G.K. and R.E.B. contributed to study design and conceptualisation. G.K, R.E.B., M.L. and C.J. collected and identified mosquitoes. S.K., J-P.P., M.L. and G.K. performed molecular identification with support from E.P. and F.B. Statistical analyses were carried out by G.K. with support from L.N. and H.M.F. The original draft was written by G.K. and reviewed by H.M.F. All authors reviewed the final manuscript.

