## Supplementary Information for "Ecology of mosquitoes in Scottish wetlands and confirmation of *Culex molestus* and hybrids in Scotland"

### Ecology of mosquitoes in Scottish wetlands and confirmation of *Culex pipiens molestus* and hybrids in Scotland – Supplementary information 1

Authors: Georgia Kirby[1,3], Rebecca E. Brown[1], Meshach Lee[1], Jean-Philippe Parvy[2], Susanne Krabbendam[1], Emilie Pondeville[2], Colin Johnston[3], Jolyon M. Medlock[3], Alexander G.C. Vaux[3], Luca Nelli[1], Francesco Baldini[1] & Heather M. Ferguson[1]

[1] School of Biodiversity, One Health and Veterinary Medicine, University of Glasgow, Glasgow, UK

[2] MRC-University of Glasgow Centre for Virus Research, Glasgow, UK

[3] Medical Entomology & Zoonoses Ecology, UK Health Security Agency, Porton Down, UK

**Table S1.** British mosquito species ecology and functional groups, adapted from Medlock *et al* 2024.

| Functional group | Oviposition site | Overwintering stage | Preferred host type | Generations | Species |
|---|---|---|---|---|---|
| **1a** | Water (vegetation) | Larvae | Mammals | Univoltine | *Cq. richiardii* |
| **1b** | Water | Larvae | Mammals | Multivoltine | *An. claviger* |
| **1c** | Water | Adult | Mammals | Multivoltine | *An. maculipennis* s.l.<br>*Cs. annulata*<br>*Cs. subochrea* |

| | | | | | |
|---|---|---|---|---|---|
| **1d** | Water | Adult | Birds | Multivoltine | *Cx. pipiens* s.l. |
| **2a** | Land | Eggs | Mammals | Univoltine | *Ae. annulipes*<br>*Ae. cantans*<br>*Ae. punctor*<br>*Ae. leucomelas** |
| **2b** | Land | Eggs | Mammals | Multivoltine | *Ae. cinereus*<br>*Ae. caspius*<br>*Ae. detritus*<br>*Ae. dorsalis** |
| **2c** | Land (tree holes) | Eggs (larvae) | Mammals | Multivoltine | *Ae. geniculatus*<br>*An. plumbeus* |
| **2d** | Land | Larvae (eggs) | Birds | Univoltine | *Cs. morsitans* |

*Species found in the present study that had not previously been classified due to rarity in Britain.

**Table S2.** List of sampling sites and associated wetland habitats.

| Site name | Coordinates | Wetland habitat(s) | Number of adult traps | Number of aquatic habitats sampled | Nearest weather station (miles from site) |
|---|---|---|---|---|---|
| Broubster Leans RSPB | 58.519573, -3.655115 | Wet grassland | 2 | 7 | Strathy East (12.3) |
| Caerlaverock NNR | 54.97132, -3.501884 | Saltmarsh, reedbeds | 2 | 4 | Dumfries, Crichton Royal No 2 (6.1) |
| Findhorn Bay | 57.638778, -3.579944 | Saltmarsh | 2 | 5 | Kinloss (0.8) |
| Gailes Marsh SWT | 55.586333, -4.656946 | Pond | 1 | 1 | Darvel, Hillview (14.7) |
| Knockan Crag NNR | 58.031583, -5.072472 | Blanket bog | 2 | 5 | Achiltibuie No 2 (9.6) |
| Knockshinnoch Lagoons SWT | 55.397753, -4.19001 | Reedbeds | 1 | 3 | Forrest Lodge, Burnhead (18.4) |
| Loch Ard Forest | 56.168986, -4.382 | Wet woodland | 1 | 4 | Gartocharn, Portnellan Farm (11.3) |
| Loch Fleet NNR | 57.937972, -4.071667 | Saltmarsh, wet grassland | 2 | 5 | Tain Range (9.2) |
| Loch Leven NNR | 56.191325, -3.403406 | Reedbeds | 1 | 0 | Auchtermuchty, Rossie (10.8) |
| Loch of Kinnordy RSPB | 56.67268, -3.043799 | Reedbeds, wet woodland | 2 | 6 | Braemar No 2 (26.1) |
| Montrose Basin SWT | 56.699444, -2.488721 | Saltmarsh | 1 | 1 | Fettercairn, Glensaugh No 2 (13.2) |
| Oldhall Pond SWT | 55.591914, -4.638779 | Pond | 1 | 4 | Darvel, Hillview (13.9) |

| | | | | | |
|---|---|---|---|---|---|
| Possil Marsh SWT | 55.903908, -4.263889 | Reedbeds, wet woodland | 2 | 7 | Glasgow, Bishopton (10.4) |
| RSPB Insh Marshes | 57.076333, -4.016 | Reedbeds | 1 | 2 | Dalwhinnie No 2 |
| RSPB Loch Leven | 56.178615, -3.356729 | Wet grassland, reedbeds | 2 | 2 | Auchtermuchty, Rossie (9.9) |
| RSPB Loch Lomond | 56.054985, -4.511525 | Wet grassland | 2 | 7 | Gartocharn, Portnellan Farm (2.2) |
| RSPB Nigg Bay | 57.733111, -4.005528 | Wet grassland, saltmarsh | 2 | 6 | Dunrobin Castle Gardens (17) |
| Shewalton Wood SWT | 55.583861, -4.635778 | Wet woodland | 1 | 13 | Darvel, Hillview (13.9) |
| Shian Wood SWT | 56.520889, -5.410417 | Saltmarsh, wet woodland | 2 | 3 | Dunstaffnage (4.9) |
| Tentsmuir NNR | 56.426333, -2.872139 | Reedbeds, pond | 2 | 2 | Leuchars (3.4) |
| The Saltings | 55.92394, -4.464363 | Saltmarsh | 1 | 1 | Glasgow, Bishopton (2.9) |
| WWT Caerlaverock | 54.976611, -3.483964 | Reedbeds, pond | 2 | 11 | Dumfries, Crichton Royal No 2 (6.2) |

**Table S3.** Mosquito abundance and sampling effort by site. Trapping night is defined as the total number of nights for which a single trap at the site was run throughout the sampling period, and is therefore doubled if two traps were present at the site. Larval dips is defined as the total number of dips taken from aquatic habitats at the site throughout the sampling period.

| Site name | Number of trapping nights | Number of adult mosquitoes collected | Total larval dips | Number of mosquito larvae collected |
|---|---|---|---|---|
| Loch of Kinnordy RSPB | 12 | 653 | 72 | 29 |
| WWT Caerlaverock | 16 | 203 | 109 | 228 |
| Possil Marsh SWT | 10 | 192 | 98 | 93 |
| Shewalton Wood SWT | 7 | 173 | 107 | 18 |
| Findhorn Bay | 12 | 155 | 45 | 15 |
| Caerlaverock NNR | 16 | 135 | 29 | 5 |
| Broubster Leans RSPB | 4 | 99 | 57 | 55 |
| The Saltings | 7 | 54 | 23 | 0 |
| Oldhall Pond SWT | 7 | 51 | 41 | 0 |
| Montrose Basin SWT | 6 | 47 | 27 | 2 |
| Tentsmuir NNR | 8 | 46 | 24 | 112 |
| RSPB Loch Lomond | 12 | 45 | 105 | 155 |
| Knockshinnoch Lagoons SWT | 5 | 31 | 29 | 119 |
| Loch Ard Forest | 6 | 15 | 42 | 5 |
| Loch Fleet NNR | 12 | 11 | 69 | 3 |
| RSPB Insh Marshes | 6 | 10 | 30 | 17 |

| | | | | |
|---|---|---|---|---|
| Gailes Marsh SWT | 2 | 8 | 6 | 0 |
| Loch Leven NNR | 2 | 7 | 0 | 0 |
| Shian Wood SWT | 8 | 6 | 42 | 0 |
| RSPB Loch Leven | 8 | 5 | 36 | 0 |
| RSPB Nigg Bay | 10 | 5 | 93 | 4 |
| Knockan Crag NNR | 8 | 0 | 39 | 0 |

**Table S4.** Definitions of aquatic habitat categories.

| Habitat type | Description | Number of representative habitats |
|---|---|---|
| Reedbed | The water/ waterlogged area surrounding a collection of reeds (*Phragmites* spp.). | 9 |
| Ditch | A constructed narrow channel that may be permanently or occasionally filled with rainwater. | 18 |
| Pool | A small, discrete, natural body of water with a maximum area of $2m^2$. Water may be permanent or transient. Includes puddles. | 35 |
| Pond | A discrete permanent body of fresh still water with an area between $2m^2$ and 2 $ha^2$. | 10 |
| Marsh | Waterlogged areas in wetlands dominated by grasses and sedges. | 16 |
| Bog | Waterlogged areas in peatlands dominated by mosses. | 3 |
| Artificial | Any rain-filled artificial container made of plastic, metal, concrete or clay. | 8 |
| Exposed | Habitats in open areas without shade from sunlight. | 54 |
| Partially shaded | Habitats in areas with patchy shelter. Some parts of the habitat are sometimes exposed to sunlight. | 26 |
| Shaded | Habitats in completely sheltered areas which are never exposed to sunlight. | 19 |

#### Text S1: Additional notes on mosquito species classification based on morphology

##### Adults

The morphologically cryptic species *Aedes cinereus* and *Aedes geminus, Culiseta morsitans* and *Culiseta litorea,* and members of the *Anopheles maculipennis* species complex were not distinguished, and are referred to as *Aedes cinereus, Culiseta morsitans* and *Anopheles maculipennis* s.l. respectively. Furthermore, several *Culiseta* subgenus *Culiseta* specimens were present without tarsi, preventing distinction between *Culiseta annulata* and *Culiseta alaskaensis*. These specimens are referred to as '*Culiseta annulata*', with the caveat that this may include some *Culiseta alaskaensis* specimens.

##### Larvae

Identification was based solely on morphology, and therefore cryptic species (*Culex pipiens* s.l./ *Culex torrentium*, and *Culiseta annulata*/ *Culiseta alaskaensis*/ *Culiseta subochrea*) were not distinguished.

#### Text S2: Molecular identification of *Culex* specimens

For all intact adult *Cx. pipiens* s.l./ *Cx. torrentium* specimens (n = 198), DNA was extracted from legs manually pulverised into 20µl of squishing buffer (10mM Tris-HCl, 1mM EDTA, 25mM NaCl) and 200g/ml proteinase K. This solution was incubated at 37ºC for 20 minutes and then inactivated by incubation at 95ºC for two minutes. The resulting DNA concentrations were checked using a Nanodrop.

PCR to identify *Culex* biotypes followed the method described in [1]. PCR products were visualised on a 2% agarose gel with SYBR Safe (Invitrogen). The size of the amplified product was approximately 200 bp for *Cx. p. pipiens* and 250 bp for *Cx. p. molestus*. Hybrids yielded two bands, one at 200 bp and one at 250 bp. Specimens that did not amplify were subjected to a second PCR cycle to confirm whether they were *Cx. torrentium*, following the method described in [2] with the PCR cycle conditions adapted to the following: three minutes at 95ºC, followed by 40 cycles of 95 ºC (30 seconds), 55ºC (30 seconds), 72ºC (40 seconds), then ten minutes at 72ºC. PCR products were again visualised on a 2% agarose gel stained with SYBR Safe. *Culex pipiens* s.l. yielded a band approximately 610 bp in size while *Cx. torrentium* bands measured approximately 416 bp. Positive *Cx. p. pipiens, Cx. p. molestus* and *Cx. torrentium* controls were obtained from colonies at the MRC-University of Glasgow Centre for Virus Research, Glasgow [3].

**Table S5**. Summary of terms tested in models. * denotes interaction between two variables. Where possible, an interaction between wetland type and previous rainfall was tested in adult abundance models to account for the possibility that the suitability of different wetland types may be mediated by the volume of rainfall. Rainfall with a three-week lag was chosen, as exploratory analysis indicated that the effect of rainfall on abundance is most evident at this interval.

| Model | Response variable | Response unit | Fixed effects | Random effects | Distribution |
|---|---|---|---|---|---|
| 1 | Total adult mosquito abundance | Number of adult mosquitoes per trap night | Latitude, longitude, wetland type, mean temperature during trapping period, total rainfall during trapping period, mean temperature lag$^{-2}$, total rainfall lag$^{-2}$, temperature lag$^{-3}$, total rainfall lag$^{-3}$, temperature lag$^{-4}$, total rainfall lag$^{-4}$, wetland type * total rainfall lag$^{-3}$ | Date of collection, site | Negative binomial |
| 2 | *Culex pipiens* s.l. adult abundance | Number of adult *Cx. pipiens* s.l. per trap night | Latitude, longitude, wetland type, mean temperature during trapping period, total rainfall during trapping period, mean temperature lag$^{-2}$, total rainfall lag$^{-2}$, temperature lag$^{-3}$, total rainfall lag$^{-3}$, temperature lag$^{-4}$, total rainfall lag$^{-4}$, wetland type * total rainfall lag$^{-3}$ | Date of collection, site | Negative binomial |
| 3 | *Anopheles claviger* adult abundance | Number of adult *An. claviger* per trap night | Latitude, longitude, wetland type, mean temperature during trapping period, total rainfall during trapping period, mean temperature lag$^{-2}$, total rainfall lag$^{-2}$, temperature lag$^{-3}$, total rainfall lag$^{-3}$, temperature lag$^{-4}$, total rainfall lag$^{-4}$, wetland type * total rainfall lag$^{-3}$ | Date of collection, site | Negative binomial |
| 4 | *Aedes detritus* adult abundance | Number of adult *Ae. detritus* per trap night | Latitude, longitude, mean temperature during trapping period, total rainfall during trapping period, mean temperature lag$^{-2}$, total rainfall lag$^{-2}$, temperature lag$^{-3}$, total rainfall lag$^{-3}$, temperature lag$^{-4}$, total rainfall lag$^{-4}$ | Date of collection, site | Poisson |
| 5 | Probability of finding *Culex pipiens* s.l/ | Presence/ absence of *Cx. pipiens* s.l/ *Cx. torrentium* larvae | Latitude, longitude, mean ambient temperature, ambient temperature lag$^{-2}$, total rainfall lag$^{-2}$, ambient | Date of collection, site | Binomial |

| | | | | | |
|---|---|---|---|---|---|
| | *Culex torrentium* larvae | | temperature lag$^{-3}$, total rainfall lag$^{-3}$, exposure, vegetation cover | | |
| 6 | Probability of finding *Culiseta annulata* larvae | Presence/ absence of *Cs. annulata* larvae | Latitude, longitude, mean ambient temperature, mean temperature lag$^{-2}$, total rainfall lag$^{-2}$, ambient temperature lag$^{-3}$, total rainfall lag$^{-3}$, exposure, vegetation cover | Date of collection, site | Binomial |
| 7 | Probability of finding *Culex pipiens* s.l/ *Culex torrentium* larvae (hydrochemical variables subset) | Presence/ absence of *Cx. pipiens* s.l/ *Cx. torrentium* larvae | Latitude, longitude, mean ambient temperature, ambient temperature lag$^{-2}$, total rainfall lag$^{-2}$, ambient temperature lag$^{-3}$, total rainfall lag$^{-3}$, exposure, vegetation cover, water temperature, pH | Date of collection, site | Binomial |
| 8 | Probability of finding *Culiseta annulata* larvae (hydrochemical variables subset) | Presence/ absence of *Cs. annulata* larvae | Latitude, longitude, mean ambient temperature, ambient temperature lag$^{-2}$, total rainfall lag$^{-2}$, ambient temperature lag$^{-3}$, total rainfall lag$^{-3}$, exposure, vegetation cover, depth, water temperature, pH, conductivity | Date of collection, site | Binomial |
| 9 | *Culex pipiens* s.l/ *Culex torrentium* larval density | Number of *Cx. pipiens* s.l/ *Cx. torrentium* larvae per dip | Ambient temperature lag$^{-2}$, ambient temperature lag$^{-3}$, exposure, vegetation cover | Date of collection, site | Negative binomial |
| 10 | *Culiseta annulata* larval density | Number of *Cs. annulata* larvae per dip | Latitude, longitude, ambient temperature lag$^{-2}$, ambient temperature lag$^{-3}$, exposure, vegetation cover | Date of collection, site | Negative binomial |

#### Text S3: Summary of microclimatic conditions during study period

Relative to the 1991-2020 period (data obtained from the Met Office), mean and maximum temperatures across Scotland in May were typical but minimum temperatures were anomalously high, reaching 2.5ºC higher than normal in the western and northern Highlands. May rainfall was lower than average, with areas of the west Highlands experiencing under 33% of average monthly rainfall. June was hotter than average, with maximum temperatures across most of the country up to 3.5ºC higher, particularly in the west. Rainfall in June was largely normal. Temperatures in July were mostly normal but were up to 1.5ºC cooler than average in the south and east. July rainfall was higher than normal, particularly in the south where it reached 175% of the average monthly rainfall. Microclimatic variables in August were mostly normal but rainfall was around 75% of the normal volume in the south west and minimum temperatures across the country were up to 1.5ºC higher than normal. September was highly anomalous, with mean and maximum temperatures up to 2.5ºC higher than normal, particularly in the south and west, and rainfall up to 200% higher than average in areas of the south.

**Table S6.** Microclimatic variables at each site during the sampling period. Mean values include ± 95% confidence interval.

| Site name | Mean temperature (ºC) | Total rainfall (mm) |
|---|---|---|
| Broubster Leans RSPB | 13.14 ± 0.47 | 361.95 |
| Caerlaverock NNR | 14.98 ± 0.42 | 483.71 |
| Findhorn Bay | 14.47 ± 0.49 | 276.97 |
| Gailes Marsh SWT | 14.42* ± 0.47 | 325.2* |
| Knockan Crag NNR | 14.06* ± 0.51 | 303.13* |
| Knockshinnoch Lagoons SWT | 13.72 ± 0.43 | 812.8 |
| Loch Ard Forest | 14.78 ± 0.39 | 553.28 |
| Loch Fleet NNR | 13.46 ± 0.45 | 304.35 |
| Loch Leven NNR | 14.1 ± 0.41 | 306.55 |

| | | |
|---|---|---|
| Loch of Kinnordy RSPB | 12.4 ± 0.43 | 343 |
| Montrose Basin SWT | 12.84 ± 0.44 | 444.97 |
| Oldhall Pond SWT | 14.42* ± 0.47 | 325.2* |
| Possil Marsh SWT | 14.81 ± 0.39 | 467.4 |
| RSPB Insh Marshes | 12.55 ± 0.44 | 406 |
| RSPB Loch Leven | 14.1 ± 0.41 | 306.55 |
| RSPB Loch Lomond | 14.78 ± 0.39 | 553.28 |
| RSPB Nigg Bay | 13.67 ± 0.43 | 372.18 |
| Shewalton Wood SWT | 14.42* ± 0.47 | 325.2* |
| Shian Wood SWT | 14.91 ± 0.38 | 593.6 |
| Tentsmuir NNR | 14.17 ± 0.42 | 285.67 |
| The Saltings | 14.81 ± 0.39 | 467.4 |
| WWT Caerlaverock | 14.98 ± 0.42 | 483.71 |

*missing data from September

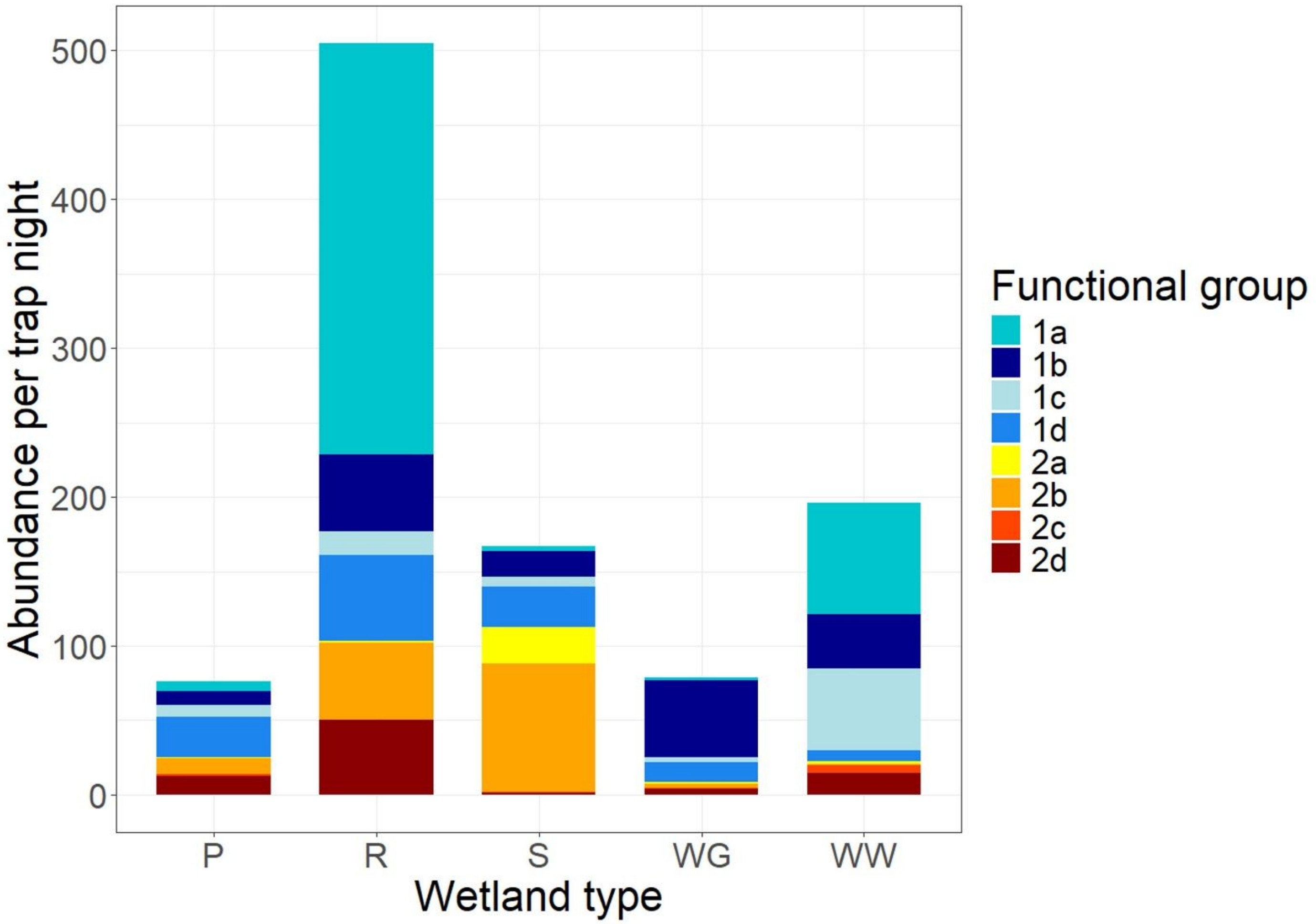

**Figure S1**. Total abundance of adult mosquito functional groups by wetland type. Abbreviations: P = Pond, R = Reedbed, S = Saltmarsh, WG = Wet Grassland, WW = Wet Woodland. Functional groups are based on the categories defined in [4]. Here, 1a (water-ovipositing mammalophilic univoltine species that overwinter as larvae) (n = 689) represents *Coquillettidia richiardii* (100%); 1b (water-ovipositing mammalophilic multivoltine species that overwinter as larvae) (n = 333) represents *Anopheles claviger* (100%); 1c (water-ovipositing mammalophilic multivoltine species that overwinter as adults) (n = 171) represents *Culiseta annulata* (95.3%), *Anopheles maculipennis* s.l. (4.1%) and *Culiseta subochrea* (0.6%); 1d (water-ovipositing ornithophilic multivoltine species that overwinter as adults) (n = 257) represents *Culex pipiens* s.l. (100%); 2a (land-ovipositing mammalophilic univoltine species that overwinter as eggs) (n = 60) represents *Aedes leucomelas* (83.3%) *Aedes punctor* (11.7%), *Aedes cantans* (3.3%) and *Aedes annulipes* (1.7%); 2b (land-ovipositing mammalophilic multivoltine species that overwinter as eggs) (n = 466) represents *Aedes detritus* (62.4%), *Aedes caspius* (21.1%), *Aedes cinereus* (8.9%) and *Aedes dorsalis* (7.6%); 2c (tree hole-ovipositing mammalophilic multivoltine species that overwinter as eggs or larvae) (n = 16) represents *Anopheles plumbeus* (93.8%) and *Aedes geniculatus* (6.2%); 2d (land-ovipositing ornithophilic univoltine species that overwinter as larvae or eggs) (n = 122) represents *Culiseta morsitans* (100%).

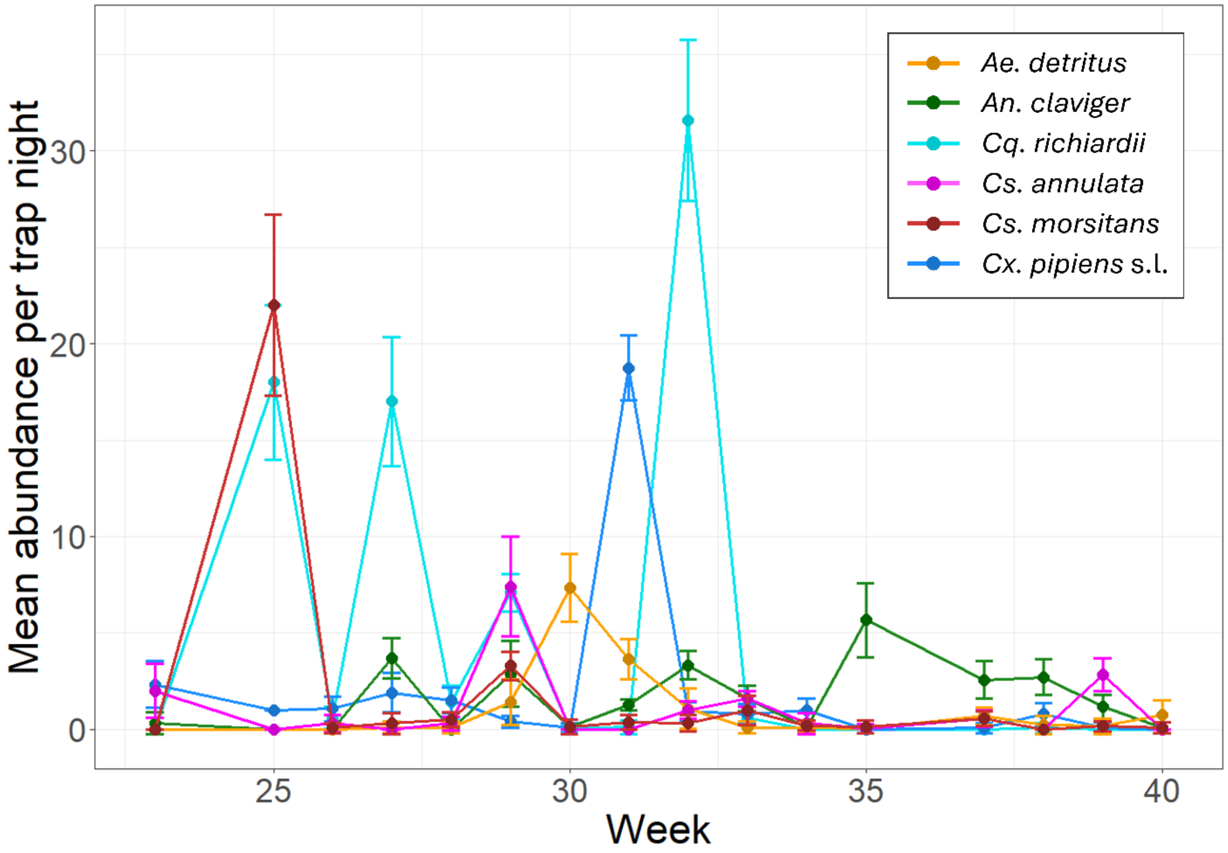

**Figure S2.** Mean weekly abundance of selected adult mosquito species at wetland sites between June and October 2023. Error bars indicate standard error.

**Table S7**. Summary of statistical significance of terms tested in adult abundance models. Chi-square ($\chi^2$) and p values are shown for each variable as obtained through likelihood ratio testing. * denotes interaction between two variables.

| Variable | $\chi^2$ | p-value |
|---|---|---|
| **Total mosquito abundance** | | |
| Latitude | 0.224 | 0.636 |
| Longitude | 3.176 | 0.075 |
| Wetland type | 2.788 | 0.594 |
| Mean temperature during trapping period | 5.344 | **0.021** |
| Total rainfall during trapping period | 0.829 | 0.363 |
| Mean temperature during trapping period $lag^{-2}$ | 0.508 | 0.476 |
| Total rainfall during trapping period $lag^{-2}$ | 0.489 | 0.484 |
| Mean temperature during trapping period $lag^{-3}$ | 0.03 | 0.862 |
| Total rainfall during trapping period $lag^{-3}$ | 6.304 | **0.012** |
| Mean temperature during trapping period $lag^{-4}$ | 1.137 | 0.286 |
| Total rainfall during trapping period $lag^{-4}$ | 1.385 | 0.239 |
| Wetland type * total rainfall during trapping period $lag^{-3}$ | 3.396 | 0.494 |
| ***Culex pipiens* s.l. abundance** | | |
| Latitude | 4.551 | **0.033** |
| Longitude | 3.261 | 0.071 |
| Wetland type | 4.918 | 0.296 |
| Mean temperature during trapping period | 3.941 | **0.047** |
| Total rainfall during trapping period | 0.152 | 0.697 |
| Mean temperature during trapping period $lag^{-2}$ | 0.001 | 0.981 |
| Total rainfall during trapping period $lag^{-2}$ | 0.013 | 0.909 |
| Mean temperature during trapping period $lag^{-3}$ | 2.122 | 0.145 |
| Total rainfall during trapping period $lag^{-3}$ | 1.248 | 0.264 |
| Mean temperature during trapping period $lag^{-4}$ | 1.616 | 0.204 |
| Total rainfall during trapping period $lag^{-4}$ | 0.001 | 0.992 |

| | | |
|---|---|---|
| Wetland type * total rainfall during trapping period lag$^{-3}$ | 6.936 | 0.139 |
| ***Anopheles claviger abundance*** | | |
| Latitude | 0.508 | 0.476 |
| Longitude | 5.628 | **0.018** |
| Wetland type | 1.681 | 0.794 |
| Mean temperature during trapping period | 0.125 | 0.724 |
| Total rainfall during trapping period | 4.017 | **0.045** |
| Mean temperature during trapping period lag$^{-2}$ | 2.902 | 0.088 |
| Total rainfall during trapping period lag$^{-2}$ | 0.694 | 0.405 |
| Mean temperature during trapping period lag$^{-3}$ | 0.966 | 0.326 |
| Total rainfall during trapping period lag$^{-3}$ | 7.327 | **0.007** |
| Mean temperature during trapping period lag$^{-4}$ | 3.286 | 0.07 |
| Total rainfall during trapping period lag$^{-4}$ | 1.534 | 0.215 |
| Wetland type * total rainfall during trapping period lag$^{-3}$ | 8.24 | 0.083 |
| ***Aedes detritus abundance*** | | |
| Latitude | 2.44 | 0.118 |
| Longitude | 2.765 | 0.096 |
| Mean temperature during trapping period | 1.843 | 0.175 |
| Total rainfall during trapping period | 0.251 | 0.617 |
| Mean temperature during trapping period lag$^{-2}$ | 1.21 | 0.271 |
| Total rainfall during trapping period lag$^{-2}$ | 0.254 | 0.614 |
| Mean temperature during trapping period lag$^{-3}$ | 1.339 | 0.247 |
| Total rainfall during trapping period lag$^{-3}$ | 1.105 | 0.293 |
| Mean temperature during trapping period lag$^{-4}$ | 2.036 | 0.154 |
| Total rainfall during trapping period lag$^{-4}$ | 14.263 | <0.000 |

**Table S8.** Summary of mean habitat characteristics of collected larval species. Mean values include ± 95% confidence interval.

| Species | Salinity (ppt) | Water temperature (ºC) | Depth (cm) | Exposure | pH | Conductivity (µS/cm) | Vegetation cover (%) |
|---|---|---|---|---|---|---|---|
| *Culex pipiens* s.l./ *Culex torrentium* | 0.1 ± 0.06 | 17.74 ± 1.21 | 8.27 ± 2.8 | Exposed | 6.83 ± 0.18 | 0.35 ± 0.14 | 42.63 ± 15.47 |
| *Culiseta annulata* | 0.16 ± 0.07 | 16.38 ± 0.83 | 13.1 ± 7.11 | Shaded | 6.86 ± 0.14 | 0.49 ± 0.16 | 62.68 ± 11.91 |
| *Anopheles claviger* | 0.23 ± 0.11 | 15.72 ± 1.34 | 10.26 ± 5.05 | Exposed | 6.92 ± 0.25 | 0.69 ± 0.2 | 54.23 ± 17.75 |
| *Culiseta morsitans* | 0.03 ± 0.04 | 14.12 ± 0.8 | 8.56 ± 2.15 | Partially shaded | 6.55 ± 0.21 | 0.22 ± 0.13 | 65.4 ± 34.52 |
| *Aedes detritus/ Aedes leucomelas* | 15 ± 0.49 | 16.67 ± 6.31 | 9.87 ± 5.93 | Exposed | 6.89 ± 0.37 | 19.47 ± 1.03 | 43.3 ± 51.03 |

**Table S9**. Summary of statistical significance of terms tested in larval presence and density models. Chi-square ($\chi^2$) and p values are shown for each variable as obtained through likelihood ratio testing. * denotes interaction between two variables.

| Variable | $\chi^2$ | p-value |
|---|---|---|
| **Probability of finding *Culex pipiens* s.l./ *Culex torrentium* larvae** | | |
| Latitude | 0.303 | 0.582 |
| Longitude | 0.065 | 0.8 |
| Mean ambient temperature | 1.476 | 0.224 |
| Ambient temperature lag$^{-2}$ | 3.366 | 0.067 |
| Total rainfall lag$^{-2}$ | 0.781 | 0.377 |
| Ambient temperature lag$^{-3}$ | 0.021 | 0.885 |
| Total rainfall lag$^{-3}$ | 0.169 | 0.681 |
| Exposure | 4.959 | 0.084 |
| Vegetation cover | 3.313 | 0.069 |

| | | |
|---|---|---|
| **Probability of finding *Culiseta annulata* larvae** | | |
| Latitude | 2.469 | 0.116 |
| Longitude | 2.397 | 0.122 |
| Mean ambient temperature | 0.247 | 0.619 |
| Ambient temperature $lag^{-2}$ | 0.063 | 0.802 |
| Total rainfall $lag^{-2}$ | 0.405 | 0.525 |
| Ambient temperature $lag^{-3}$ | 0.197 | 0.657 |
| Total rainfall $lag^{-3}$ | 0.5 | 0.479 |
| Exposure | 13.245 | **0.001** |
| Vegetation cover | 8.716 | **0.003** |
| **Probability of finding *Culex pipiens* s.l/ *Culex torrentium* larvae (hydrochemical variables subset)** | | |
| Latitude | 1.667 | 0.197 |
| Longitude | 0.005 | 0.944 |
| Mean ambient temperature | 1.652 | 0.199 |
| Ambient temperature $lag^{-2}$ | 0.065 | 0.8 |
| Total rainfall $lag^{-2}$ | 1.982 | 0.159 |
| Ambient temperature $lag^{-3}$ | 0.28 | 0.597 |
| Total rainfall $lag^{-3}$ | 0.697 | 0.404 |
| Exposure | 5.547 | 0.062 |
| Vegetation cover | 4.423 | 0.035 |
| Water temperature | 8.183 | **0.004** |
| pH | 0.011 | 0.916 |
| **Probability of finding *Culiseta annulata* larvae (hydrochemical variables subset)** | | |
| Latitude | 3.228 | 0.072 |
| Longitude | 1.799 | 0.18 |
| Mean ambient temperature | 0.615 | 0.433 |
| Ambient temperature $lag^{-2}$ | 0.587 | 0.444 |

| | | |
|---|---|---|
| Total rainfall lag$^{-2}$ | 0.741 | 0.389 |
| Ambient temperature lag$^{-3}$ | 0.622 | 0.43 |
| Total rainfall lag$^{-3}$ | 0.015 | 0.903 |
| Exposure | 8.756 | **0.013** |
| Vegetation cover | 4.294 | **0.038** |
| Water temperature | 2.029 | 0.154 |
| pH | 0.021 | 0.886 |
| Conductivity | 4.265 | **0.039** |
| ***Culex pipiens* s.l/ *Culex torrentium* larval density** | | |
| Ambient temperature lag$^{-2}$ | 2.691 | 0.101 |
| Ambient temperature lag$^{-3}$ | 0.017 | 0.897 |
| Exposure | 5.265 | 0.072 |
| Vegetation cover | 2.602 | 0.107 |
| ***Culiseta annulata* larval density** | | |
| Latitude | 2.9 | 0.089 |
| Longitude | 1.035 | 0.309 |
| Ambient temperature lag$^{-2}$ | 0.294 | 0.588 |
| Ambient temperature lag$^{-3}$ | 0.114 | 0.736 |
| Exposure | 11.113 | **0.004** |
| Vegetation cover | 3.429 | 0.064 |

1 Bahnck, C.M. and Fonseca, D.M. (2006) Rapid assay to identify the two genetic forms of *Culex* (*Culex*) *pipiens* L. (Diptera: Culicidae) and hybrid populations. *Am J Trop Med Hyg* 75, 251-255

2 Smith, J.L. and Fonseca, D.M. (2004) Rapid assays for identification of members of the Culex (Culex) pipiens complex, their hybrids, and other sibling species (Diptera: culicidae). *Am J Trop Med Hyg* 70, 339-345

3 Parvy, J.-P., Brass, D.P., Preston, B., Carmichael, R., Kerrigan, D., Kirby, G., Brown, R., White, S.M., Ferguson, H.M., Pondeville, E. (In Press) Field Colonisation and Temperature-Driven Variation in Life-History Traits of the West Nile Virus Vector Culex pipiens at Its Northern European Limit. *Parasites & Vectors*
4 Medlock, J.M.*, et al.* (2024) Mosquito diversity and abundance in English wetlands – empirical evidence to guide predictions for wetland suitability for mosquitoes (Diptera: Culicidae). *Journal of the European Mosquito Control Association* 42, 3-29
